# Pest control strategies after first signs of invasive black rat: case study in the Community of Madrid

**DOI:** 10.1101/2021.01.19.427231

**Authors:** Azucena Bermejo-Nogales, José M. Navas

## Abstract

Rodents are animals that provoke special concern in the rural and urban areas as may cause potential damage in facilities and infrastructures as well as social alarm. The control of rodent populations is based on prevention and on what it has been called an “Integrated Pest Management” strategy. The most important species affected by this strategy are brown rat (*Rattus norvegicus*), black rat (*Rattus rattus L*.) and mouse (*Mus musculus*). In the present work, our original objective was to monitor changes in status or range of rodent species in the Community of Madrid (Spain). We conducted in coordination with the professional organization of Pest Control Operators “ANECPLA” a procedure designed to obtain faecal samples in locations with rodenticide treatment. Determination of pest specie was based on cytochrome b (*cytb*) sequences and phylogenetic analysis. We received samples from a variety of locations in which a rodenticide treatment was applied due to infestation or to citizen notice. While we recorded a number of data about the distribution of brown rat the presence of black rat was unexpectedly discovered. The detection of this species implies changes in pest control strategies to improve the results in the application of rodenticides available in the market.

**Key Message:** Rodent pest control is a key issue but little is known about Spanish populations. We aimed to know how many rat species there are in Central Spain, specifically in the Community of Madrid. We found for the first time the presence of black rat and a new wild strain of brown rat in this area. These findings have change the strategies of rodent treatment and stresses the importance of population studies for a better control.

## Introduction

Rodents are a potential cause of disease for human beings, domestic animals and wildlife and they may cause potential damage in facilities and infrastructures as well as social alarm. Therefore, surveillance and control of rodent populations are key aspects for public health. There is international consensus on the need to adopt prevention and control strategies based on an “Integrated Pest Management” strategy, as reflected in the European Union (EU) Directive 2009/128/EC (EC, 2009) on the sustainable use of pesticides and in its amendment Directive 2019/782/EC (EC, 2019) regarding the establishment of harmonised risk indicators. In the European strategy, special attention is paid to some rodent species, in particular brown rat (*Rattus norvegicus*), black rat (*Rattus rattus L*.) and mouse (*Mus musculus*). The most important challenges in this strategy are those related with the management of pests favoured by climate change and with the evolution of pesticide resistance. At the same time, the strategy faces the complexity of effective pest management strategies in a context of pesticide availability reduction. Additional identified challenges in the strategy are decreasing research, increasing scarcity of human expertise, lack of knowledge transfer into practice, and some communication gaps within and between countries. This strategy includes as one of the main themes the importance of favouring preventive measures for sustainable pest management. Therefore, the preliminary and continuous evaluation of hazards and risks as well as preventive / control measurements on the environment should be aimed at minimizing the use of chemical substances.

A fundamental aspect in the studies about pest control is this related with the evaluation of the presence of invasive species. It is essential to generate detailed information about the new locations of invasive species and compile data about their behaviour and habits in order to design a successful pest control strategy. Notwithstanding the foregoing, the general opinion establishes the need of fundamental research as regards to control of invasive species, especially invasive black rat. In Spain, this information does not currently exist and the present study is part of the government strategy for monitoring rodent pest species.

Normally, studies related with pest control are based on the collection of living or dead individuals in the field in order to obtain tissues or blood samples that are used to monitor rodenticide activity or to perform genomic analysis (Goulois et al. 2017; Grandemange et al 2009; Obiegala et al. 2019). However, the improvement of DNA extraction techniques allows nowadays the use of faecal samples for this kind of analyses. Faecal samples are extremely common and easy to collect in comparison to fresh tissues, increasing in this way the number of samples available for the study without the need for sampling by trapping or managing dead animals (Meerburg et al. 2014).

Our main goal in this first step was to monitor important changes in the status or range of rat species in the Community of Madrid. We conducted in coordination with “ANECPLA”, the Spanish professional organization of Pest Control Operators (PCOs), a procedure designed to obtain faecal samples in locations of the Community of Madrid in which an intensive rodenticide treatment was applied due to infestation or to citizen notice. Rodent species were determined by analysis of the *cytochrome b* (*cytb*) gene in DNA extracted from faecal samples. This cytochrome is as a marker of variability among species (Irwin et al. 1991) and was previously used to differentiate species in the Rattini tribe (Pages et al. 2010). The *cytb* sequences were blasted into the Genebank database and aligned for phylogenetic analysis. The existence of brown rat was confirmed while strikingly and unexpectedly black rat presence was observed.

## Material and Methods

### Survey and sampling

A survey of rodent populations with the final aim of detecting possible resistances to rodenticides has been carried out at the Autonomous Community of Madrid (Spain) concentrating on those places where a number of rodent sightings were reported. The survey was designed in coordination with “ANECPLA”, the Spanish professional organization of Pest Control Operators (PCOs) to inquire resistance in the brown rat, as was the only rat specie that was expected. For this survey, a questionnaire with the characteristics of the location of the faecal samples were associated with a unique barcode. This barcode was used to track the samples for the correct traceability during sample determination. The questionnaire was included in a kit consisted on material for the collection of stool samples (a sterile cryogenic vial with the unique barcode, gloves, spatula and instructions to collect the sample). The kit was prepared in our laboratory and distributed to PCOs. Initially 50 kits were supplied.

### Homogenization of droppings and DNA extraction

One complete faecal dropping (200 mg approximately) was homogenized for each location. For comparative and control purposes, we used two controls. The first one consisted of fresh tail tissue (final 0.5 cm) obtained from a death brown rat supplied by PCOs. The second one was one sample (10^7^ cells) of the brown rat hepatoma cell line H4IIE, obtained from the European Collection of Animal Cell Cultures (ECACC) (Wiltshire, UK). H4IIE were cultured in MEM Eagle (EMEM) media (Ref.:12–125, Lonza, Barcelona, Spain) with 1% non-essential amino acids, 1% L-Glutamine, 1% Penicilline/streptomycin and 10% foetal bovine serum (Bermejo-Nogales et al. 2017).

Each sample (controls and faecal samples) was homogenized twice in a 2 ml eppendorf safe-lock containing 1 ml of sterile TE and one steel ball of 5 mm (Werfen Ref.: BAI5-1000) for 2 min at 30 frequency with the Tissue Lyser II (QUIAGEN, Hilden, Germany). Thereafter a first centrifugation step was performed to discard excessive non-degradable matter at 10,000 x *g* for 30 sec (standard rotor F45–24–11, Eppendorf centrifuge 5415R). The supernatant was collected and added to 900 µl of fresh PBS and vortexed. A second centrifugation was carried out at 4,000 x *g* for 15 min to obtain the pellet for DNA extraction. The pellet was re-suspended in 200 µl of BT1 buffer with 25 µl of proteinase K and incubated overnight at 56°C for DNA isolation using a SPEEDTOOLS DNA extraction kit (BIOTOOLS, Madrid, Spain) according to the manufacturer’s instructions. DNA yield of all samples was 10-500 ng/µl with 260 and 280 nm UV absorbance ratios (A260/280) of 1.9-2.1.

### Cytochrome b gene amplification by PCR

Rodent species were identified by analysis of the *cytb* gene in DNA extracted from faecal samples. Partial DNA sequences coding for *cytb* were PCR amplified (a heating cycle at 94 °C for 3 minutes followed by 35 cycles at 94 °C for 30 s, 50 °C for 30 s and 72 °C for 90 s) with the DreamTaq Hot Start Greeen PCR Master Mix (Thermofisher scientific, Waltham, MA, US) in a final volume of 50 μl. Forward 5’-TCTCCATTTCTGGTTTACAAGAC-3’ and reverse 5’-AACAATGACATGAAAAATCATCGTT-3’ primers were previously designed from the available literature (Pages et al. 2010). Amplified PCR products were gel-extracted and purified with the Illustra Exo ProStar kit (Sigma Aldrich, Madrid, Spain) according to the manufacturer’s instructions. Products were sequenced by the deoxy chain termination method (ABI PRISM dRhodamine terminator cycle sequencing kit, Perkin–Elmer, Wellesley, MA, USA).

### Sequence analysis: BLAST and phylogenetic tools

A BLAST search strategy was used to corroborate the identity of amplified *cytb* products that share amino acid identity with genes of either brown rat or black rat (**Table 1**). Multiple sequence alignments were carried out with ClustalW software. The phylogenetic tree was constructed on the basis of nucleotide differences with the Maximum Likelihood (ML) algorithm in MEGA7 (Kumar et al. 2016). Sequences of nine representative Rattini species found in other world regions (Pages et al. 2010) was used in the analysis. Mouse *cytb* sequence was used as out-group. Reliability of the tree was assessed by bootstrapping, using 1000 bootstrap replications.

**Table 1.** E-values and Identity percentage (brackets values) of cytochrome b sequences from faecal samples with reference sequences from brown rat and black rat available in public databases.

| Faecal sample<br>Bar code | Length<br>(bp) | Brown rat (KY356141) | Black rat<br>(FJ355927) | Location / Zip code |
| --- | --- | --- | --- | --- |
| Y2602309<sup>+</sup> | 7 | - | - | Madrid/ 28001 |
| Y2602325<sup>+</sup> | 99 | - | - | Alcobendas/ 28100 |
| Y2602289<sup>+</sup> | 216 | 3e-105 (99.53) | - | Rivas Vaciamadrid/<br>28522 |
| Y2602258<sup>+</sup> | 45 | 8e-11 (100) | - | Alcobendas/ 28100 |
| Y2602283 | 833 | 0.0 (100) | - | Alcobendas/ 28100 |
| Y2602320 | 879 | 0.0 (100) | - | Coslada/ 28821 |
| Y2602315 | 977 | 0.0 (100) | - | Coslada/ 28821 |
| Y2602282 | 943 | 0.0 (100) | - | Las Rozas/ 28230 |
| Y2602296 | 882 | 0.0 (100) | - | Tres Cantos/ 28760 |
| Y2602292 | 943 | 0.0 (100) | - | Madrid/ 28001 |
| Y2602298 | 679 | 0.0 (100) | - | Madrid-Pinar del rey/<br>28033 |
| Y2602293 | 937 | 0.0 (100) | - | Madrid/ 28001 |
| Y2602319 | 315 | 0.0 (100) | - | Madrid-Barajas/ 28042 |
| Y2602294 | 913 | 0.0 (100) | - | Madrid-Numancia/ 28038 |
| Y2602322 | 864 | 0.0 (99.77) | - | Arganda del Rey/ 28500 |
| Y2602321 | 686 | 0.0 (99.56); 0.0 (100)* | - | Madrid/ 28001 |
| Y2602340 | 787 | - | 0.0 (100) | Madrid-Peñagrande/<br>28035 |
| Y2602311 | 969 | - | 0.0 (100) | Madrid-Valdefuentes/<br>28055 |
| Y2602303 | 932 | - | 0.0 (100) | Alcobendas/ 28100 |
| H4IIE | 878 | 0.0 (99.54); 0.0 (100)* | - |  |
| TAIL | 906 | 0.0 (99.89) | - | Colmenar Viejo/ 28770 |

<sup>+</sup> Low quality sequence.
\* E-value and Identity percentage with the strain Wild/Tku (DQ673917).

## Results

### Survey and sampling

Eight companies gave a positive response, collecting 19 samples between May and June 2018. In order to maintain confidentiality, the location was only referenced with the corresponding postal code. Samples showed a wide distribution inside the Community of Madrid area (**Figure 1A**). All companies indicated the presence of brown rat as possible specie for the samples.

**Figure 1.**
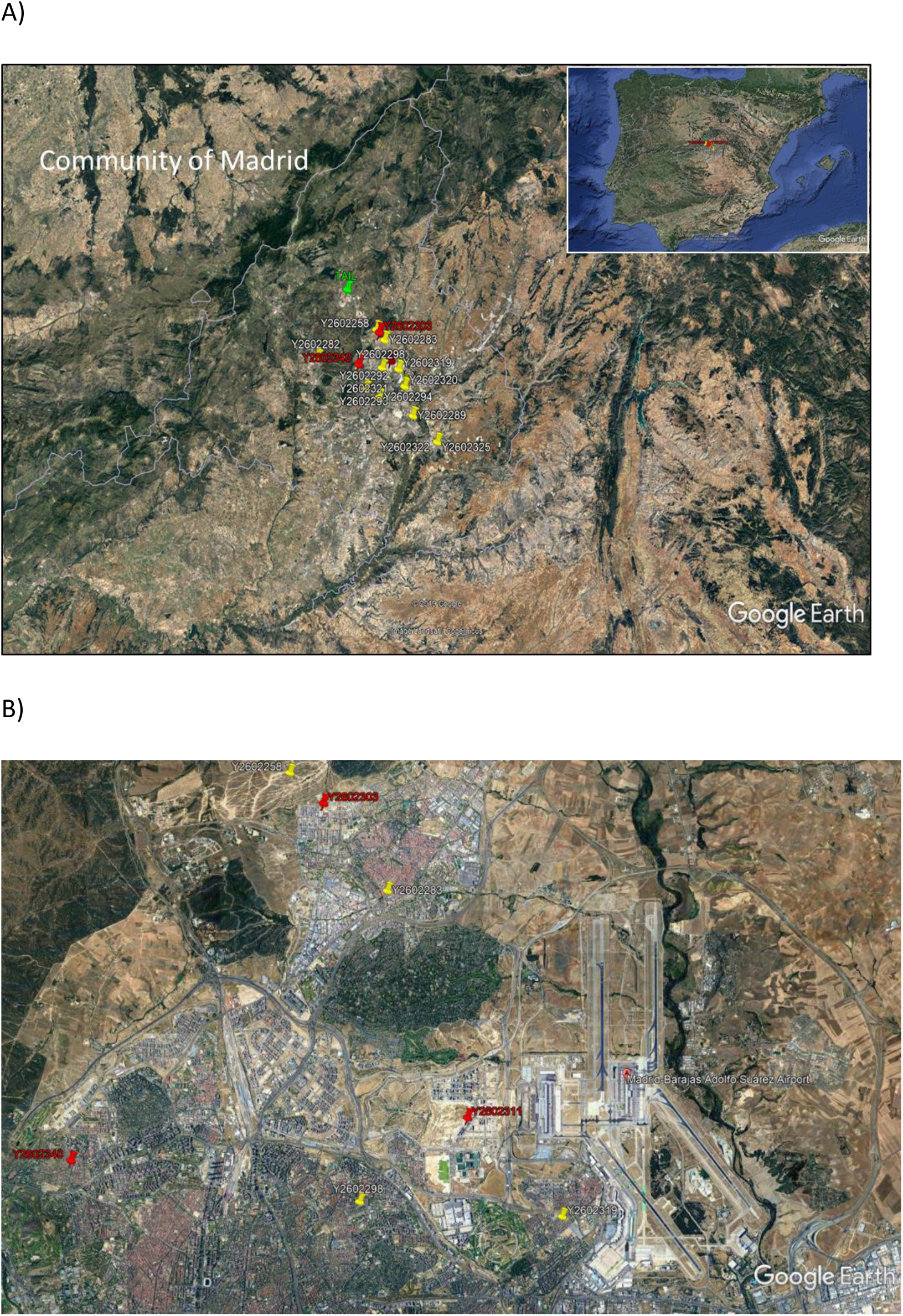
Google Earth view with the distribution map of received samples. A) Community of Madrid. B) Detail of black rat locations. Control tail sample (green), brown rat (yellow) and black rat (red).

### Samples and sequence analyses

A total of 19 DNA sequences obtained from faecal samples and the two control samples (tail and brown rat hepatoma cell line H4IIE) were analysed. The 21 sequences were from different quality, four traces with short-medium contiguous read length and 17 traces with long contiguous read length. We obtained sequences with a median length of 915 nucleotides and a range of 99 to 1012 nucleotides. The phylogenetic tree was constructed based on high quality sequences that were previously revised and curated. **Figure 2** shows Maximum Likelihood and Bootstrap trees corresponding to 28 *cytb* sequences analysed that encompass sequences obtained from15 faecal samples, two control sequences (one from brown rat tail and one from the H4IIE rat hepatoma cell line). All positions with less than 95% site coverage were eliminated. That is, fewer than 5% alignment gaps, missing data, and ambiguous bases were allowed at any position. There were a total of 626 positions in the final dataset. The phylogenetic analysis of *cytb* sequences from faecal samples groups 12 sequences as brown rat and three sequences as black rat reinforcing the evidence of the presence of samples coming from this species among those collected and analysed.

**Figure 2.**
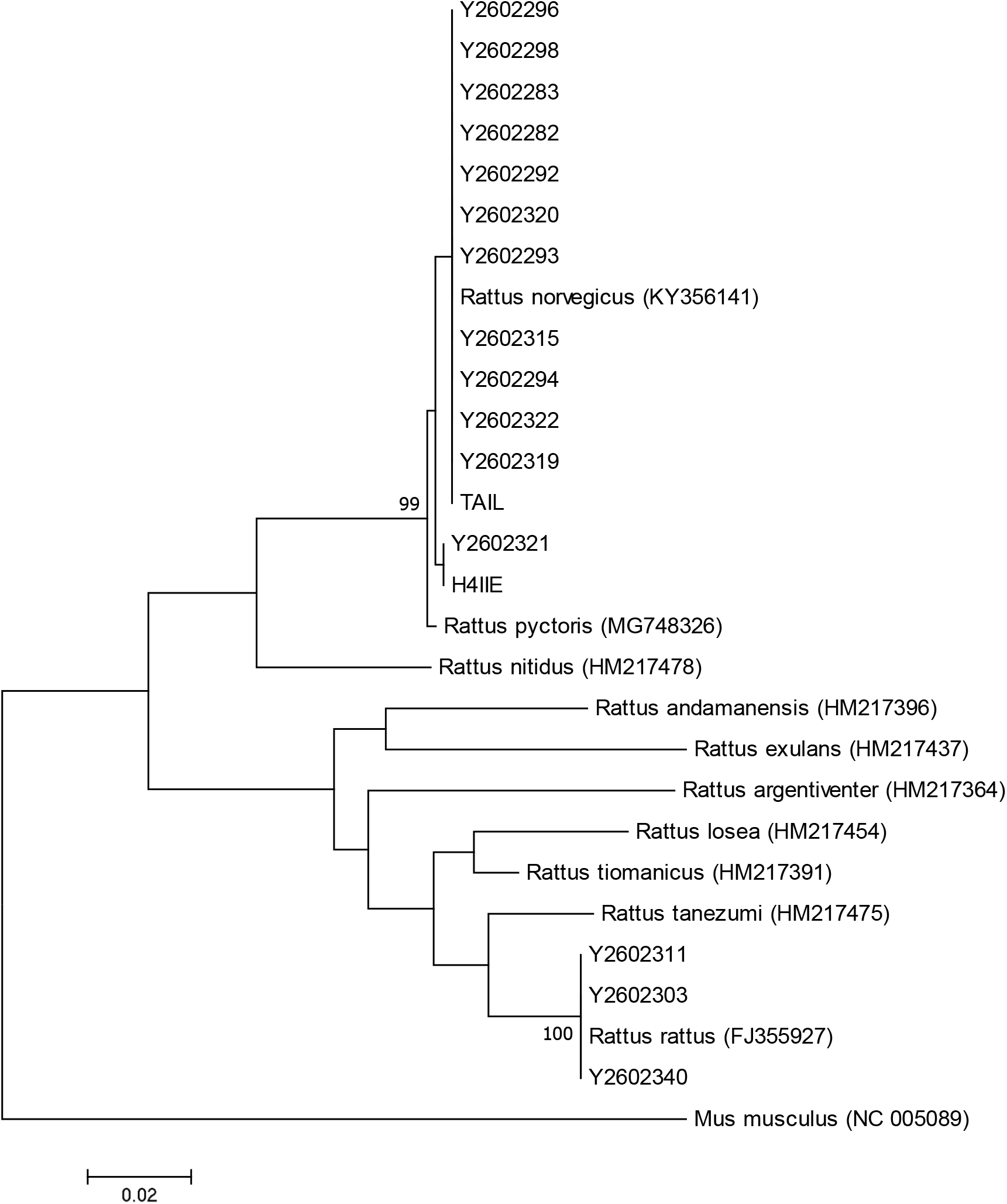
Molecular Phylogenetic analysis by Maximum Likelihood method of 27 sequences that encompasses 15 DNA faecal samples, two control sequences and different rat species. Bar code of each faecal sample is indicated for traceability. Accession number of for cytb of each rat species is indicated in brackets.

BLAST analysis was also done in order to reveal possible species definition (**Table 1**). All sequences identified as brown rat had an identity percentage higher than 99% with the reference brown rat sequence (KY356141). Nonetheless, the sequence corresponding to the faecal sample Y2602321 has a 99.56% with this reference sequence but has a 100% identity with the *Rattus norvegicus* strain Wild/Tku sequence (DQ673917). The phylogenetic tree also places this sequence within the H4IIE branch of brown rat. After this procedure, the four low quality sequences (previously discarded for phylogenetic analysis) were analysed by BLAST. Two sequences did not yield any results but the two medium-length sequences could be identified as brown rat.

Regarding location, the three samples identified as black rat are located in the vicinity of Madrid-Barajas Adolfo Suarez airport in an area with strong anthropogenic disturbances (**Figure 1B** and **Table 1**).

## Discussion

Today, the need for a correct identification of pest specie is more urgent than ever since biodiversity in many areas is being altered rapidly by the ongoing global change. Rodents are among the pest species that produce a highest social alarm. Therefore, there is a need to control their populations and the use of rodenticides is considered nowadays as one of the most appropriate ways to control them. The use of these chemicals is regulated under various legislations.

The arboreal black rat is originated from Asia and was spread to Europe and throughout the world. This species inhabits in warm areas being displaced by the brown rat in cooler regions and urban areas. In the Iberian Peninsula and the Canary Islands, black rat has a wide distribution pattern in rural and urban zones, but is mainly located in coastal or temperate areas where the climate is more permissive (López et al. 2013). This initial distribution pattern has led to the general assumption among PCOs that the rodent species responsible for common infestations in the Iberian plateau (in the centre of the Iberian Peninsula) is the brown rat. Thus, the three black rat samples received in our laboratory were identified by PCOs as brown rat. Our results, however, demonstrate for the first time the presence of black rat in the Community of Madrid, in the very centre of the Iberian plateau. Several factors could favour the presence of this invasive species in this location. The arrival of black rat in this area could have a certain causal relationship with the international freight transport activity in Madrid that is concentrated at the airport (Lopez-Escolano et al. 2019). Indeed, the black rat was detected in areas with high anthropogenic disturbance that are more prone to be occupied by opportunistic mammals (Cavada et al. 2019). In addition, probably the progressive increase in temperature detected during the past years (Vicente-Serrano et al. 2017) has also favoured the settlement of this temperate species.

A particular issue revealed by the present study is this related with the possible presence of new strains of brown rat in Madrid. Multiple strains of brown rat have been identified across the world (Schlick et al. 2006). In the present study, we have identified one sample with 100% identity with the wild strain Wild/Tku that it is originally from Tokyo. Therefore, the possibility of settlement of new variants of brown rat may not be discarded and further research needs to be done.

All the above has important consequences for the control of possible rat invasions at the Community of Madrid, but this has also implications at a broad perspective. In general, rodent control is performed with the use of anticoagulant rodenticides. First generation anticoagulant rodenticides require several days of feeding to be fully active and their lethal dose 50 (LD50, the dose of rodenticide ingested by the animal to cause 50% of lethality) is higher than those of second generation anticoagulant rodenticides that are fully active just after 24 h of feeding. Studies on the toxicological effects of these substances have been carried out for brown rat and mice, and evidenced important sensitivity differences among species (Wheeler et al. 2019). Therefore, the confident differentiation of species is a remarkable result for the appropriate management of rodents since it allows the application of the appropriate rodenticide treatment (in terms of concentration of rodenticide in bait, amount of bait, and placement of bait). For example, in the case of an infestation, PCOs could treat with brodifacoum (a second generation anticoagulant rodenticide) at 0.4 mg/kg body weight in the case of brown rat while this treatment has to reach 0.8 mg/kg body weight in the case of black rat and up to 2 mg/kg body weight in the case of mice (Wheeler et al. 2019). A specific species-bait concentration for the treatment of each pest should be part of a more sustainable use of pesticides favoured by the scientific knowledge. In line with this tendency to a more sustainable use of pesticides, the current EU rodenticide regulation (Regulation 2016/1179 amending Regulation 1272/2008 (EC, 2016)) establishes a reduction in the concentration of rodenticides in bait to nearly half (<30 mg/kg bait) of the previous one e (50 mg/kg bait) (Frankova et al. 2019) in order to minimize harmful consequences for wildlife (Quinn et al. 2019; Seljetun et al 2019; Wiens et al 2019).

In conclusion, in invasive species control programs, it is of high importance to identify first and foremost each rodent species or variant for an appropriate assessment of pest risks. The present study demonstrates for the first time the presence of black rat in the Community of Madrid and urges towards an appropriate management of mouse, black rat and brown rat control considering differences in sensitivity among species against anticoagulant rodenticides.

## Declarations

### Availability of data and material

The data will be available upon request

### Author’s contributions

ABN and JMN conceived the study and contributed to the text. ABN perform all the analysis. JMN led the EG17-017 project.

